# Genetic characterisation of candidate ecdysteroid kinases in *Drosophila melanogaster*

**DOI:** 10.1101/2024.01.22.576657

**Authors:** Jack L. Scanlan, Charles Robin

**Affiliations:** School of BioSciences, The University of Melbourne, Parkville Campus, Melbourne, Victoria, 3010, Australia

**Keywords:** ecdysone, 20-hydroxyecdysone 3-dehydroecdysone trachea, UAS/GAL4 RNAi CRISPR, steroid

## Abstract

Ecdysteroids are major hormones in insects and control moulting, growth, reproduction, physiology, and behaviour. The biosynthesis of ecdysteroids such as 20-hydroxyecdysone (20E) from dietary sterols is well characterised, but ecdysteroid catabolism is poorly understood. Ecdysteroid kinases (EcKs) mediate the reversible phosphorylation of ecdysteroids, which has been implicated in ecdysteroid recycling during embryogenesis and reproduction in various insects. However, to date only two EcK-encoding genes have been identified, in the silkworm *Bombyx mori* and the mosquito *Anopheles gambiae*. Previously, we identified two ecdysteroid kinase-like (EcKL) genes—*Wallflower* (*Wall*) and *Pinkman* (*pkm*)—in the model fruit fly *Drosophila melanogaster* that are orthologs of the ecdysteroid 22-kinase gene *BmEc22K*. Here, using gene knockdown, knockout and misexpression, we explore *Wall* and *pkm*’s possible functions and genetically test the hypothesis that they encode EcKs. *Wall* and *pkm* null mutants are viable and fertile, suggesting they are not essential for development or reproduction, whereas phenotypes arising from RNAi and somatic CRISPR appear to derive from off-target effects or other artefacts. However, misexpression of *Wall* results in dramatic phenotypes, including developmental arrest, and defects in trachea, cuticle and pigmentation. *Wall* misexpression fails to phenocopy irreversible ecdysteroid catabolism through misexpression of *Cyp18a1*, suggesting Wall does not directly inactivate 20E. Additionally, *Wall* misexpression phenotypes are not attenuated in *Cyp18a1* mutants, strongly suggesting Wall is not an ecdysteroid 26-kinase. We hypothesise that the substrate of Wall in this misexpression experiment and possibly generally is an unknown, atypical ecdysteroid that plays essential roles in *Drosophila* development, and may highlight aspects of insect endocrinology that are as-yet uncharacterised. We also provide preliminary evidence that *CG5644* encodes an ecdysteroid 22-kinase conserved across Diptera.

## Introduction

Ecdysteroids are polyhydroxylated steroids that control numerous aspects of insect biology, most notably moulting, growth and reproduction (Riddiford 1993). Twenty-hydroxyecdysone (20E) is responsible for most of the functions attributed to ecdysteroids, although its precursor ecdysone (E) may also have specific developmental roles (Ono 2014; Beckstead et al. 2007). The nuclear receptor heterodimer EcR/USP and the G protein-coupled receptor DopEcR are thought to modulate most genomic and non-genomic responses to ecdysteroids (Ishimoto et al. 2012; Baker et al. 2000; Wang et al. 2000), although the nuclear receptor DHR38 may also be involved in ecdysteroid signalling (Baker et al. 2003). The biosynthesis of ecdysteroids from dietary sterols involves the highly conserved “Halloween” enzymes (Rewitz et al. 2006, 2007) and the tight transcriptional regulation of Halloween genes is essential for controlling insect growth and development (Kannangara et al. 2021; Yamanaka et al. 2013). However, relatively little is known about ecdysteroid inactivation through catabolism and how this is regulated. Ecdysteroid inactivation can modulate ecdysteroid signalling by reducing levels of active hormone and/or producing inactive “storage” forms that can be recycled into active hormones. This can play important roles in reproduction, where maternal- or mate-contributed storage hormones can influence early development or reproductive physiology, respectively (Sonobe & Ito 2009; Peng et al. 2022; Crocker & Hunter 2018). It can also function in development, by shaping the pulse-like titres of 20E required for developmental transitions (Rewitz et al. 2010). Ecdysteroid recycling might also be responsible for the mid-pupal pulse of 20E in some brachyceran dipterans (Scanlan et al. 2023; Ohtaki 1981), although this does not appear to be the case in *Drosophila melanogaster* (Zhang et al. 2023).

Ecdysteroid catabolic reactions can be broadly classified into modifications and conjugations. Modifications involve changes to various hydroxyl groups on the ecdysteroid nucleus. 26-hydroxylation occurs via Cyp18a1 (Guittard et al. 2011) and can result in ecdysteroids that may be functional in some insects but not others (Baker et al. 2000; Kaplanis et al. 1973). 26-hydroxylated ecdysteroids can be further oxidised by Cyp18a1 to 26-ecdysonoic acids, which is an irreversible process and widespread across taxa (Rewitz et al. 2010; Li et al. 2014). 3-oxidation and 3-epimerisation also occur through ecdysteroid 3-oxidases and 3-alpha-reductases (Takeuchi et al. 2001; Sun et al. 2012), although the former modification can be reversed through the action of 3-beta-reductases (Chen et al. 1996; Sakurai et al. 1989). Conjugations involve the addition of a bulky polar or non-polar moiety that is thought to broadly limit the binding affinity to ecdysteroid receptors (Baker et al. 2000; Makka et al. 2002). Conjugated moieties include glucose (Thompson et al. 1987), fatty acids (Hoffmann et al. 1985) and acetate (Modde et al. 1984), although the most common is phosphate.

Phosphorylation reactions can occur at four potentially hydroxylated carbons on the ecdysteroid nucleus—C-2, C-3, C-22 and C-26—but the prevalence of each seems to vary broadly across both substrates and taxa (reviewed in Lafont et al. 2012). Some ecdysteroid-phosphates can be hydrolysed *in vivo* to active ecdysteroids through the action of phosphatases (Yamada & Sonobe 2003; Davies et al. 2007; Isaac, Sweeney, et al. 1983) but others may be non-hydrolysable, and therefore terminal, catabolites (Lagueux et al. 1984; Sonobe & Yamada 2004).

The most well-defined function for ecdysteroid-phosphates in insect biology is ovary– embryo recycling: phosphate conjugates are formed in, or transported to, the oocyte and hydrolysed to active hormones after fertilisation. *Bombyx mori* (Lepidoptera: Bombycidae) recycle 22-phosphates of 20E and some precursors in its eggs (Sonobe & Ito 2009), while the related moth *Manduca sexta* (Lepidoptera: Sphingidae) recycles 26-hydroxy-E 26-phosphate (Feldlaufer et al. 1987). Two locust species, *Schistocerca gregaria* and *Locusta migratoria* (Orthoptera: Acrididae), both recycle ecdysteroid 22-phosphates (Isaac, Sweeney, et al. 1983; Isaac, Rose, et al. 1983; Lagueux et al. 1984). Hydrolysis of ecdysteroid-phosphates seems to control embryonic diapause in *B. mori* (Matsushima et al. 2019; Yamada et al. 2005) and may do the same in locusts (Tawfik et al. 2002). Various ecdysteroid-phosphates are also produced during later life stages of lepidopterans and orthopterans but their functions are unclear (Beydon et al. 1987; Gibson et al. 1984; Modde et al. 1984; Lozano et al. 1989).

In Diptera, ecdysteroid phosphorylation saw comparatively limited study until recently. In the mosquito *Anopheles gambiae* (Diptera: Culicidae), 3-dehydro-20E 22-phosphate is a male sexual gift that is dephosphorylated in the female reproductive tract to prevent remating (Peng et al. 2022). 20E is also converted to a 22-phosphate in virgin females of this species to modulate the hormonal response to blood-feeding (Peng et al. 2022). In *Musca domestica* (Diptera: Muscidae), ecdysteroid 2-phosphorylation can be induced by a synthetic ecdysteroid agonist but its physiological function remains unclear (Williams et al. 2002). In *D. melanogaster*, four ecdysteroid-phosphates have been identified: 3-dehydro-E 2-phosphate and 3-epi-20E 3-phosphate in larvae (Hilton 2004; Sommé-Martin et al. 1988a); E 22-phosphate in adult ovaries (Grau et al. 1995; Pis et al. 1995); and 20,26-dihydroxy-E 26-phosphate in the S2 cell line, due to the necessity of a phosphatase pre-treatment when characterising Cyp18a1 (Guittard et al. 2011). Possible roles for these metabolites in development or reproduction have yet to be identified.

Ecdysteroid-phosphate conjugates are produced by ecdysteroid kinases (EcKs); reverse genetic analyses of EcK genes could be a powerful method to understand the functions of ecdysteroid-phosphates in insect biology. To date, only two EcKs have been biochemically and genetically characterised: BmEc22K, which is responsible for ecdysteroid 22-phosphate production in the *B. mori* ovary (Sonobe et al. 2006); and AgEcK2, which produces 20E 22-phosphate in *A. gambiae* virgin females (Peng et al. 2022). Both *BmEc22K* and *AgEcK2* belong to the ecdysteroid kinase-like (EcKL) gene family. We recently performed a comprehensive phylogenomic analysis of the EcKL family in 140 insect genomes, classifying them into 13 subfamilies. *BmEc22K* and *AgEcK2* belong to the A and D subfamilies, respectively, making other genes in these clades strong *a priori* candidates for encoding EcKs (Scanlan & Robin 2023). *D. melanogaster* contains two subfamily A genes: *CG13813*—which we name *Wallflower* (*Wall*) in this study—and *Pinkman* (*pkm*), previously named *CG1561*. Both genes have been lost 2–3 times in Diptera but are otherwise conserved as single-copy orthologs in most species (Scanlan & Robin 2023), and they are more closely related to each other than they are to any lepidopteran subfamily A EcKL (Figure S1).

*pkm* is strongly expressed in the adult eye, with some expression in the pupal CNS and fat body (Graveley et al. 2011; Leader et al. 2018; Xu et al. 2004). The Pkm protein has an N-terminal disordered domain, atypical of most EcKLs, and may physically interact with the small heat shock proteins Hsp23 and Hsp26 and an uncharacterised carbonyl reductase, CG11200 (Guruharsha et al. 2011). It has been proposed that Pkm modulates synapse formation through interactions with Hsp23 and Hsp26 (Santana et al. 2020). *pkm* also appears to be transcriptionally induced in adult oenocytes during starvation, under the control of insulin signalling (Chatterjee et al. 2014); given this, we hypothesise *pkm* has a role in the metabolic response to starvation.

*CG13813* is primarily expressed in the midgut of embryos, larvae and adult flies (Leader et al. 2018; Weiszmann et al. 2009). Despite a lack of focused study, multiple transcriptional data suggest *CG13813* has an ecdysteroid-related function. Specifically, *CG13813* has: (1) enriched expression in the 3^rd^-instar larval ring gland (RG), which contains the ecdysteroidogenic prothoracic gland (PG) cells (Ou et al. 2016); (2) temporal co-expression with *Cyp18a1* (Graveley et al. 2011; Figure S2), especially during the mid-embryonic pulse of ecdysteroids (Maróy et al. 1988); (3) strong induction by exogenous 20E in nearly all surveyed cell lines (Stoiber et al. 2016; Gauhar et al. 2009); and (4) positive regulation by the primary 20E response gene *DHR3* (Ruaud et al. 2009; Mazina et al. 2015). Given that *CG13813* is strongly expressed in S2 cells during 20E exposure and is co-expressed with *Cyp18a1* during embryogenesis, we hypothesise that *CG13813* could encode the ecdysteroid 26-kinase proposed by Guittard et al. (2011).

Here, we genetically characterise the EcKL genes *pkm* and *CG13813* (which we name *Wallflower*/*Wall*) through germline and somatic CRISPR knockouts, RNAi knockdown and UAS-GAL4 misexpression. We also test two specific hypotheses: (1) that *Wall* encodes an ecdysteroid 26-kinase, through epistasis experiments with *Cyp18a1*; and (2) that *pkm* is involved in the starvation response in adult flies. We aim to genetically identify EcKs in *D. melanogaster*, in order to shed light on their functions in this species and across insects more broadly.

## Materials and Methods

### Fly lines and husbandry

The following fly lines (BL, Bloomington line) were obtained from the Bloomington Drosophila Stock Center (BDSC): *w\**;; *Sb^1^*/TM3 *actGFP Ser^1^*(BL4534), *Tango11^1^*/CyO *actGFP* (BL36320), FM7i *actGFP*/C(1)DX *y^1^ f^1^* (BL4559), FM7j (BL6418), *y^1^ w^67c23^*;; *P{EPgy2}EY20330* (BL23106, called UAS-*Wall^EY^*herein), *w\**; *nos-GAL4*; UAS-*Cas9.P2* (BL67083) and *elav-GAL4*; UAS-*Cas9.P2*/CyO (BL67073). The following UAS-dsRNA (and control genotype) lines were obtained from the Vienna Drosophila Resource Centre (VDRC): VL60100 (‘KK control’), *w^1118^* (VL60000; ‘GD control’), *KK^Wall^* (VL104249), *GD^Wall^* (VL45409), *KK^pkm^* (VL106503), *GD^pkm1^* (VL32634) and *GD^pkm2^*(VL32635). UAS-*Cyp18a1* (Guittard et al. 2011), UAS-*Dcr2*/CyO; *tub-GAL4*/TM6B and some GAL4 driver stocks (Table S2) were a kind gift of Philip Batterham (The University of Melbourne). *w\**; *tub-GAL80^ts^*; TM2/TM6B and *y^1^ w* P{lacW}Fas2G0032 P{neoFRT}19A*/FM7c; *P{ey-FLP.N}5* were a kind gift of Michael Murray (The University of Melbourne). The strong loss-of-function *Cyp18a1^1^*allele was described in Rewitz *et al*. (2010); a *Cyp18a1^1^*/FM7i *actGFP* line was a kind gift of Michael O’Connor (University of Minnesota). For GAL4 driver lines used in this study, see Table S1; other fly lines were made by routine crossing (Supplementary Methods).

For routine stock maintenance, flies were kept on yeast-cornmeal-molasses media (‘lab media’; http://bdsc.indiana.edu/information/recipes/molassesfood.html) at 18 °C, 21 °C or 25 °C in plastic vials sealed with cotton stoppers.

### Generation of UAS-ORF lines

Cloning of ORFs into the pUASTattB vector (Bischof et al. 2007) and *D. melanogaster* transformation was as described in Scanlan et al. (2022), with the following modifications. The *Wall* ORF was isolated by PCR from the DGRC cDNA clones IP11764 and GH09153, using the primer pair CG13813_EagIF and CG13813_KpnIR (Table S2). The two ORFs of *pkm*—the full 635 aa ORF (‘*pkm^F^*’) and a truncated 430 aa ORF without the N-terminal intrinsically disordered region (‘*pkm^T^*’)—were synthesised by Integrated DNA Technologies, with *EagI* and *KpnI* restriction sites (plus six additional nucleotides to allow for efficient digestion) at the N- and C-termini, respectively. Expected amplicon sizes from recombinant plasmids in colony PCR for *Wall*, *pkm^F^* and *pkm^T^*were 1,521 bp, 2,136 bp and 1,521 bp, respectively (using primers pUASTattB_3F and pUASTattB_5R; Table S2). All correctly assembled plasmids, as verified by Sanger sequencing, were sent to TheBestGene Inc. (US) for microinjection and incorporation into the *D. melanogaster* genome at the attP40 site on chromosome 2 (BL25709).

### Generation of CRISPR-Cas9 mutant lines

The recombinant pCFD6 plasmids pCFD6-CG13813 and pCFD6-CG1561—each of which express, under the control of a UAS promoter, four gRNAs that target either the *CG13813*/*Wall* locus (Figure 1 A) or the *CG1561*/*pkm* locus (Figure 1B)—were designed *in silico* using Benchling (http://benchling.com), and cloned as described in Scanlan et al. (2022), with the following modifications: the primer pairs for the amplification of the inserts for pCFD6-CG13813 were pCFD6_CG13813_1F/R, pCFD6_CG13813_2F/R and pCFD6_CG13813_3F/R; and the primer pairs for the amplification of the inserts for pCFD6-CG1561 were pCFD6_CG1561_1F/R, pCFD6_CG1561_2F/R and pCFD6_CG1561_3F/R (Table S2). Correctly assembled plasmids, as verified by Sanger sequencing, were sent to TheBestGene Inc. for microinjection and incorporation into the *D. melanogaster* genome at the attP40 site on chromosome 2 (BL25709), to produce the homozygous fly lines UAS-*Wall^pCFD6^* and UAS-*pkm^pCFD6^*.

**Figure 1.**
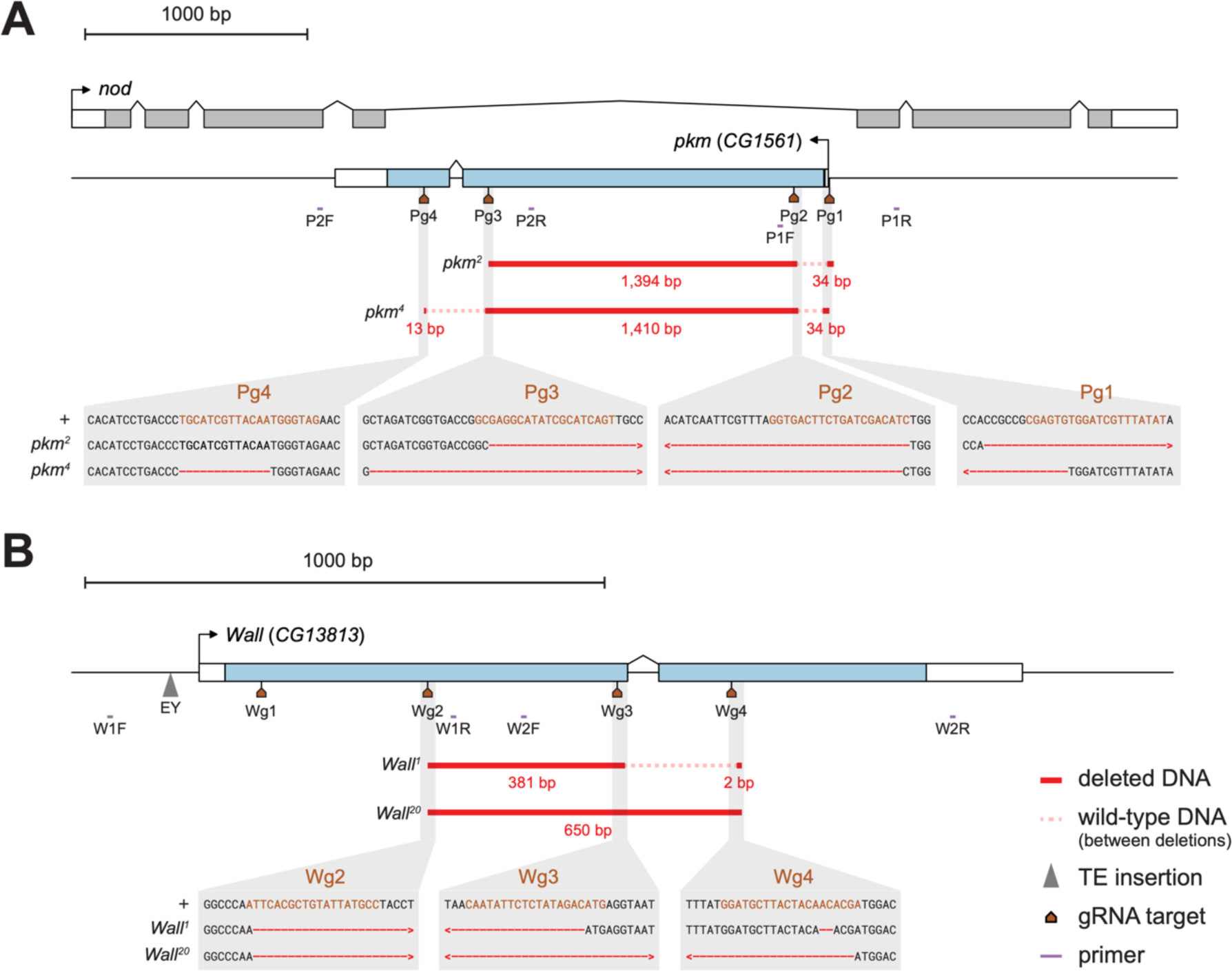
(A) The *Pinkman*/*pkm* (*CG1561*) locus on the X chromosome of *Drosophila melanogaster*, including the *pkm* gene (blue coding regions) and the *nod* gene (grey coding regions). Mapped onto the locus are the locations of gRNA target sites (Pg1–4; brown), genotyping primer-binding sites (purple), and two CRISPR-Cas9-induced deletion alleles (*pkm^2^* and *pkm^4^*; deletion sizes in red text). P1F, ΔD37_1F; P1R, ΔD37_1R; P2F, ΔD37_2F; P2R, ΔD37_2R. (B) The *Wallflower*/*Wall* (*CG13813*) locus on chromosome 3L of *Drosophila melanogaster*. Mapped onto the locus are the locations of gRNA target sites (Wg1–4; brown), genotyping primer-binding sites (purple), the *EYgp2* element *EY20330* (grey triangle), and two CRISPR-Cas9-induced deletion alleles (*Wall^1^* and *Wall^20^*; deletion sizes in red text). Neither deletion allele had lesions at the Wg1 target site. W1F, ΔD38_1F; W1R, ΔD38_1R; W2F, ΔD38_2F; W2R, ΔD38_2R. Wild-type DNA present in-between deletions is indicated with pale dashed lines. Grey boxes (bottom) are sequence-level detail of the deletions with respect to the wild-type genetic background (+). gRNA target sites are highlighted in brown, and deleted bases are red dashes; > and < symbols indicate that the deletion continues out of the frame of the highlighted sequence.

Mutagenesis of the *Wall* locus on chromosome 3L followed protocols previously described in Scanlan et al. (2022), using the crossing scheme in Figure S3A. Mutagenesis of the *pkm* locus on the X chromosome was done using the crossing scheme in Figure S3B. Single founder female flies during *pkm* mutagenesis were allowed to lay viable larvae before undergoing DNA extraction.

*Wall* PCR genotyping used the ΔD38_1F/1R and ΔD38_2F/2R primer pairs, which produce amplicons of size 698 bp and 845 bp, respectively, from wild-type chromosomes, and ΔD38_1F/2R, which produce 1,689 bp amplicons from wild-type chromosomes and amplicons between ∼700–1,500 bp in the case of deletions between any two *Wall* gRNA pairs. *pkm* PCR genotyping used the ΔD37_1F/1R and ΔD37_2F/2R primer pairs, which produce amplicons of size 693 bp and 898 bp, respectively from wild-type chromosomes, and ΔD37_2F/1R, which produce 2,757 bp amplicons from wild-type chromosomes and amplicons between ∼900–2,600 bp in the case of deletions between any two *pkm* gRNA pairs. PCR—2 min initial denaturation (95 °C), then 2 min denaturation (95 °C), 45 sec annealing (55 °C) and 1.5 min extension (72 °C) for 32 cycles, then a 5 min final extension (72 °C)—was carried out with GoTaq Green Master Mix (Promega). Deletion loci from mutant flies were amplified using the genotyping primers above with four GoTaq Green PCR reactions per line, which were combined before gel-purification to allow for the detection of early-cycle polymerase-derived errors by close inspection of the sequencing chromatogram output. Gel-purified amplicons were sequenced using the appropriate genotyping primers at AGRF.

### Germline knockout, somatic knockout, RNAi knockdown and misexpression developmental viability assays

Egg-to-adult and larval-to-adult viability assays were conducted as described in Scanlan et al. (2022), but at 25 °C or 29 °C depending on the experiment. The ‘MultinomCI’ function in the *DescTools* package in R was used to calculate confidence intervals for multinomial proportions.

Misexpression phenotyping was conducted by crossing 3–5 UAS-responder females to *GAL4* males, letting them lay in lab media vials for 24 hours, with a minimum of three replicate vials per genotype; vials were left to develop at 25 °C and were checked every 24 hours for developmental arrest phenotypes. Arrest phenotypes were fully penetrant unless otherwise noted.

### Delayed-onset misexpression assay

Three groups of 10 *act-GAL4*/CyO *actGFP* virgin females were crossed to *tub-GAL80^ts^*; UAS-*Wall^EY^* males and allowed to mate in lab media vials at 18 °C for 24 hours, then transferred to three fresh vials at 18 °C (the GAL80^ts^ restrictive temperature) and allowed to lay for 24 hours. This transfer-and-lay process was repeated every 24 hours (with fresh females added after nine days) until 19 sets of three vials containing offspring were produced, and then all the vials were shifted to 29 °C (the GAL80^ts^ permissive temperature) to complete development—each set of vials contained animals that started expressing *Wall^EY^* at a different developmental stage. Eclosing adults were genotyped based on the presence or absence of the phenotypic markers on the CyO *actGFP* chromosome. Adult genotype counts from each set of vials were analysed by the ‘binom.test’ function in R (with a Bonferroni correction for multiple tests) to determine if genotypic ratios were significantly different from the Mendelian expectation (1:1).

### Food avoidance assay

*tub-GAL80^ts^*; *da-GAL4* virgin females were crossed to *w^1118^* or UAS-*Wall^EY^* males and laid eggs on juice plates (Supplementary Methods) at 25 °C for eight hours. 1^st^-instar larvae were transferred to lab media vials (20 larvae per vial, eight vials per genotype) and kept at 29 °C. Two and three days post-hatching, the number of larvae located in the food substrate (mouthparts hidden in substrate; ‘digging’), on top of the food substrate (mouthparts out of substrate; ‘on food’) or on the sides of the vials (body completely off substrate; ‘side of vial’) was scored for each genotype. Fisher’s exact test was used to determine if there were significant differences in larval position between genotypes and/or timepoints, using the ‘fisher.test’ function in R.

### Misexpression epistasis assays

Six *Cyp18a1^1^*/FM7i *actGFP*; UAS-*Wall^pU^*virgin females or +; UAS-*Wall^pU^* virgin females were crossed to *da-GAL4* or *Mef2-GAL4* males and allowed to lay in lab media vials for 24 hours, with 10 vials per genotype. Offspring were left to develop at 25 °C for 14 days, and the number of individuals that reached pupariation, pupation (or pupariation/pupation for *da-GAL4* crosses), pharate adult differentiation and eclosion were scored. Fisher’s exact test was used to determine if there were significant differences between the developmental outcomes of offspring of each maternal genotype, using the ‘fisher.test’ function in R.

### Starvation assays

Starvation assays were conducted at 21–22 °C using the ‘wet starvation’ method developed by Storelli et al. (2019). Adult flies were collected as virgins in the first 12 hours post-eclosion, separated by sex and kept in lab media vials for 2–3 days. Empty vials were half-filled with deionised water and cellulose acetate stoppers (Flystuff, 49-102) were pushed to the bottom and saturated. Excess water was thoroughly removed and 9–10 flies were placed into each vial (six vials per sex-genotype combination), which were closed with an additional dry stopper. Flies were moved to fresh vials twice a week. Survival was scored every 24 hours until all flies were dead. Male and female flies were analysed separately due to known sex differences in starvation resistance (Jang & Lee 2015). Survival analysis was conducted with the *survival* (v2.38) and *survminer* (v0.4.7) packages in R, using a log-rank test to compare survival curves; pairwise p-values were adjusted for multiple tests using the Benjamini-Hochberg procedure.

### Additional misexpression assays

UAS-ORF responder lines for 13 EcKLs were obtained from FlyORF at the University of Zurich (Bischof et al. 2013) or the DPiM transgenic fly resource at the Bangalore Fly Resource Center (Guruharsha et al. 2014); FlyORF lines contain 3xHA-tagged ORFs, while DPiM lines contain FLAG-HA-tagged ORFs. FlyORF EcKL lines used were F002982 (*CG31300*), F002821 (*CG10560*), F002832 (*CHKov2*; ‘*CHKov2^1^*’) and F002521 (*CG9259*). DPiM lines used were 817 (*CG10562*), 854 (*CHKov2*; ‘*CHKov2^2^*’), 1332 (*JhI-26*), 2262 (*CG14314*), 2439 (*CG5644*), 2866 (*CG10514*), 3380 (*CG6830*), 3774 (*CG31102*), 3915 (*CG31087*) and 3916 (*CG31288*). *w^1118^* (VDRC stock 60000) was used as a wild-type control in the absence of true matched genetic backgrounds. *tub-GAL4*/TM3 *actGFP Ser^1^* and *phm-GAL4*/CyO *actGFP* (Guittard et al. 2011) were a kind gift of Philip Batterham (The University of Melbourne).

UAS-ORF responder (and *w^1118^*) males were crossed to *tub*-GAL4/TM3 *actGFP Ser^1^* females, which were allowed to lay on lab media food, and the offspring were left to develop at 25 °C for 14 days. Adult offspring were collected after eclosion and scored for the presence or absence of the TM3 *actGFP Ser^1^* balancer chromosome (and the TM6B *Antp^Hu^ Tb^1^* chromosome in the case of DPiM line 3774). UAS-*CG5644* males were also crossed to *phm-GAL4*/CyO *actGFP* females; adult offspring were scored for the presence or absence of the CyO *actGFP* chromosome. The ‘binom.test’ function in R was used to test deviations from expected Mendelian ratios (1:1 for all crosses except those involving DPiM line 3774, which was 1:3)—significant deviations against misexpression genotypes were considered evidence for developmental lethality due to misexpression of the EcKL ORF in question.

## Results

### *pkm* null mutants have no obvious developmental phenotypes, in contrast to somatic knockout and some RNAi knockdown animals

We explored the developmental essentiality of *pkm* through three gene disruption methods: germline CRISPR, to generate heritable null alleles; somatic CRISPR, to induce gene knockout in somatic tissues; and transgenic RNAi, to knock down mRNA levels. Through germline CRISPR mutagenesis, 18 putative deletions at the *pkm* locus were generated from 24 founder females. Two *pkm* composite deletion alleles were kept as homozygous-viable stocks and molecularly characterised: *pkm^2^*and *pkm^4^*. *pkm^2^* comprises two deletions, one of 34 bp overlapping the transcription start site and another of 1,394 bp that induces a frameshift in exon 1, while *pkm^4^* comprises three deletions, one of 34 bp that removes the transcription start site, one of 1,410 bp in-frame in exon 1, and another of 13 bp that induces a frameshift in exon 2 (Figure 1A). Both alleles delete Brenner’s motif, induce frameshifts and delete significant portions of the coding sequence, and are therefore likely strong loss-of-function (null) alleles.

The developmental essentiality of *pkm* was tested with egg-to-adult viability assays involving null mutants. In crosses between *pkm^2^*/FM7c females and +/Y males, while FM7c hemizygotes appeared much less viable than other genotypes, there were no clear differences between the proportions of other genotypes, showing that *pkm^2^*/Y hemizygotes are developmentally viable (Figure 2B).

**Figure 2.**
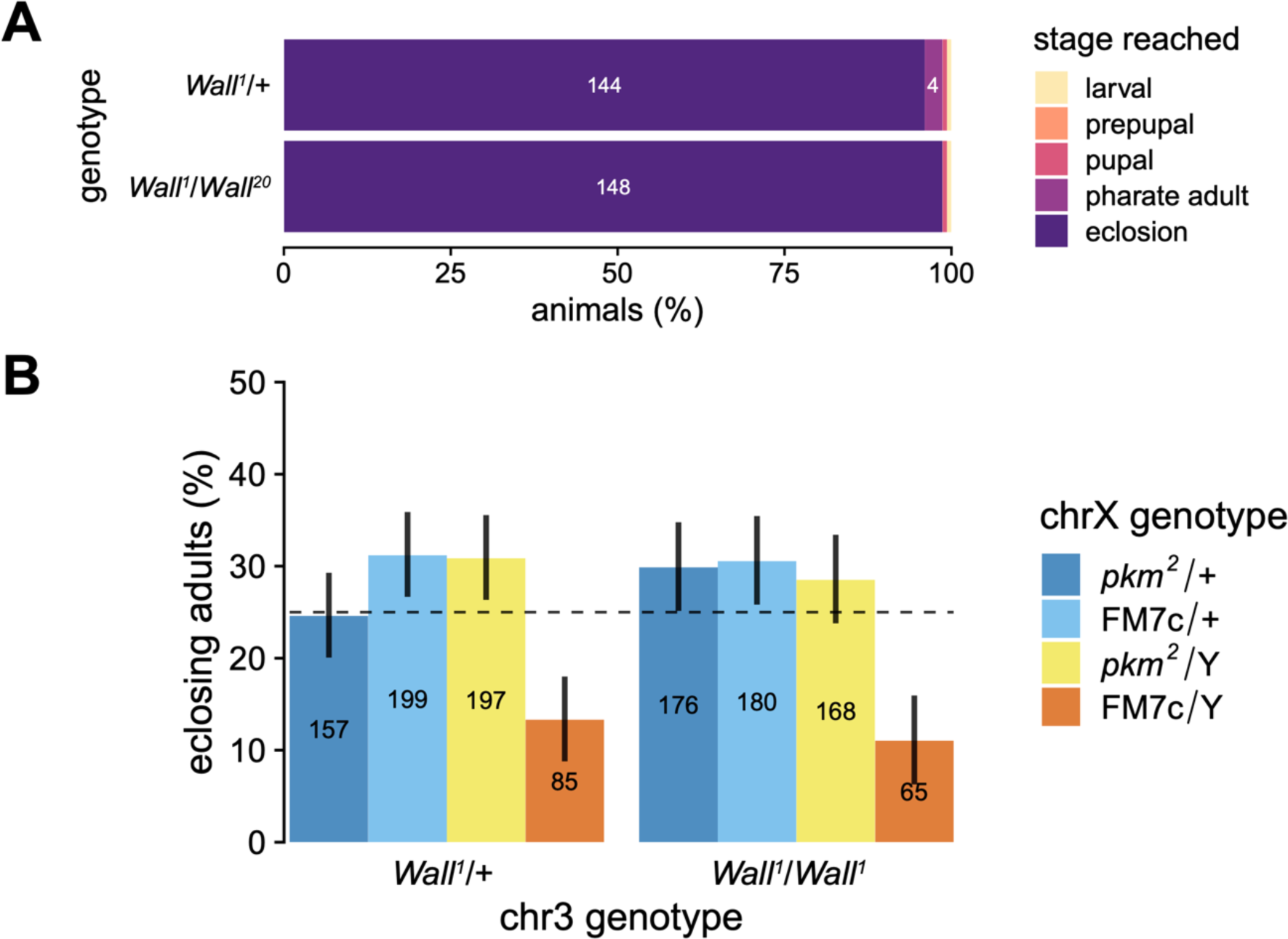
Developmental viability of *Wall* and *pkm* single and double mutant animals. (A) Larval-to-adult viability of offspring from crosses between *Wall^1^* females and wild-type (+) males (top) or *Wall^20^* males (bottom). Numbers on the bars are the number of individuals in each lethal phase category (for numbers greater than three). (B) Eclosing adult genotypic ratios of the offspring from crosses between *pkm^2^*/FM7c;; *Wall^1^* females and *Wall^1^* (left) or *w^1118^* (right) males. Bars are coloured by their chrX genotype, as determined by their sex and the presence or absence of the FM7c balancer chromosome; *pkm^2^* hemizygotes are in yellow (single mutants on left, double mutants on right). The dashed line indicates the expected 1:1:1:1 genotypic ratio if all genotypes per cross are equally developmentally viable. Numbers on the bars are the number of adults of each genotype. Error bars are 97.5% confidence intervals (95% CI adjusted for two tests) for the proportion of each genotype within each cross.

Somatic knockout of *pkm* using the ubiquitous *da-GAL4* driver resulted in a substantial number of adults falling into the food media, being unable to remove themselves and ‘drowning’; these adults were unable to be easily counted. Tipping vials upside-down before eclosion revealed that knockout adults eclosed in a 1:1 ratio with balancer adults, indicating they had no detectable pre-adult lethality (Figure S4B), but had an uncoordinated limb phenotype, effectively resulting in immobility (Figure S4C), a phenotype not seen in *pkm* null mutants. This immobility phenotype was also observed when UAS-*pkm^pCFD6^* and UAS-*Cas9* were driven with the (non-specific; see Berger et al. 2007; Casas-Tintó et al. 2017) neuronal driver *elav-GAL4*, although this was not quantified.

RNAi knockdown experiments using the strong ubiquitous *tub-GAL4* driver were previously performed at 25 °C with KK library UAS-*dsRNA* lines (Scanlan et al. 2020), but we added to this experiment by also using independently generated GD lines that target different parts of each gene’s transcript (Dietzl et al. 2007). We also performed knockdown at a higher temperature (29 °C), which increases GAL4 activity and should result in stronger knockdown (Duffy 2002; Fortier & Belote 2000), as well as at 29 °C with the addition of a UAS-*Dcr2* transgene, which can further increase knockdown efficiency (Dietzl et al. 2007).

For *pkm*, none of the three RNAi constructs produced detectable developmental arrest at 25 °C (Figure S5A), but *KK^pkm^* produced partially penetrant lethality before the adult stage at 29 °C (Figure S5B). *KK^pkm^*, *GD^pkm1^* and *GD^pkm2^* all had significant effects on developmental viability compared to controls (all p < 2.2×10^-16^, Fisher’s exact test) at 29 °C when paired with the UAS-*Dcr2* transgene, with *KK^pkm^* resulting in pupal and pharate adult lethality, and *GD^pkm1^* and *GD^pkm2^* producing a mix of pharate adult lethality and a ‘drowning’ phenotype whereby adults eclosed from the puparium but fell to the bottom of the vial and failed to escape from the semi-liquid food media (Figure S5D).

### *Wall* null mutants and somatic knockout animals have no obvious developmental phenotypes, in contrast to most RNAi knockdown animals

We also explored the developmental essentiality of *Wall* through germline CRISPR, somatic CRISPR and transgenic RNAi. Through germline CRISPR mutagenesis, 22 putative deletions at the *Wall* locus were detected by PCR from 66 founder males. Two *Wall* deletion alleles were kept as homozygous-viable stocks and molecularly characterised: *Wall^1^*and *Wall^20^*. *Wall^1^* comprises two deletions in the coding region of the genes—a 381 bp in-frame deletion in exon 1 (between sgRNAs Wg2 and Wg3), and a 2 bp frameshift deletion in exon 2 (at sgRNA Wg4). *Wall^20^* comprises a single 605 bp frameshift deletion that spans exon 1 and 2 (Figure 1B). Both alleles are predicted to significantly disrupt the function of the encoded protein, due to the deleted coding sequence (including the ATP-binding Brenner’s motif) and frameshifts, and are therefore likely also null alleles.

The developmental essentiality of *Wall* was tested with larval-to-adult viability assays involving null mutants. There were no significant differences between the developmental outcomes of *Wall^1^*/+ and *Wall^1^*/*Wall^20^*heterozygotes (p = 0.152, Fisher’s exact test; Figure 2A), with nearly all individuals successfully eclosing as viable adults. To test for possible developmental redundancy between *Wall* and *pkm*, we also conducted egg-to-adult viability assays with crosses that produced double mutant animals and compared the results to those that produced *pkm* single mutants (see above). There was no significant difference in the genotypic ratios between *Wall^1^*/*Wall^1^*and *Wall^1^*/+ crosses (p = 0.168, Fisher’s exact test; Figure 2B), demonstrating double mutants (homozygous for *Wall^1^* and hemizygous for *pkm^2^*) do not suffer any developmental viability loss compared to other genotypes.

We also noted that all generated *Wall* and *pkm* mutant stocks could be kept for multiple generations as homozygous stocks without obvious issues, suggesting neither gene is required for fertility, although this was not quantified.

Somatic knockout of *Wall* produced no detectable loss of egg-to-adult viability compared to non-gRNA-expressing animals (Figure S4A), consistent with the viability of germline null mutants. In contrast, most RNAi knockdowns of *Wall* affected developmental progression. The *KK^Wall^*RNAi construct (but not *GD^Wall^*) resulted in detectable developmental arrest before the adult stage at either 25 °C (Figure S5A) or 29 °C (Figure S5B). Both *KK^Wall^* and *GD^Wall^* had significant effects on developmental viability compared to controls (both p < 2.2×10^-16^, Fisher’s exact test) at 29 °C when paired with the UAS-*Dcr2* transgene, with *KK^Wall^*resulting in mostly larval lethality, and *GD^Wall^* resulting in both pharate adult lethality and the ‘drowning’ phenotype seen with *GD^pkm1^*and *GD^pkm2^* knockdowns (Figure S5D).

### RNAi-induced developmental arrest may be due to off-target effects, not genetic compensation

We considered our knockdown results to be suspicious, given that no developmental phenotypes were detected for *Wall* and *pkm* null mutant animals (Figure 2). However, discrepancies between null mutant and RNAi knockdown animals can be due ‘genetic compensation’, wherein gene knockout triggers a compensatory mechanism (such as the up-regulation of genes with similar functions) to restore a wild-type phenotype, while RNAi knockdown fails to trigger compensation, leading to a mutant phenotype (El-Brolosy & Stainier 2017; El-Brolosy et al. 2019). To test the hypothesis that the phenotypic discrepancy between knockdown animals and null mutant animals was due to genetic compensation in mutants, we performed *pkm* knockdown in a *pkm^2^*null mutant background, with the expectation that if genetic compensation was occurring in mutants, knockdown would fail to produce a developmental arrest phenotype, while if the knockdown phenotype was due to off-target effects, the presence of the *pkm^2^* allele would have no effect on the phenotype. Females homozygous for the *KK^pkm^* construct—and homozygous for either the *pkm^2^*allele (one of two such independently generated lines) or a wild-type allele (*pkm^+^*)—were crossed to *tub-GAL4*/TM3, *actGFP*, *Ser^1^* males, and adult genotypic ratios of the offspring were scored, with half of the knockdown individuals (males) expected to be *pkm^+^* or *pkm^2^*hemizygotes. The proportion of adult knockdown individuals between knockdown crosses with *pkm^+^* and *pkm^2^* was not significantly different (Figure S5C), with a *pkm^2^* (line 1) vs *pkm^+^* odds ratio of 1.67 (97.5% CI: 0.94, 3.0) and a *pkm^2^* (line 2) vs *pkm^+^* odds ratio of 1.02 (97.5% CI: 0.54, 1.91). This demonstrates that the presence or absence of *pkm* loss-of-function alleles does not affect the *tub>KK^pkm^* knockdown phenotype, strongly suggesting these phenotypes are due to off-target effects and not genetic compensation. A summary of all *pkm* and *Wall* knockout and knockdown phenotypes, for easy comparison, is presented in Table S3.

### Ubiquitous misexpression of *Wall*, but not *pkm*, causes developmental arrest

Without obvious requirements for *Wall* or *pkm* during development, we sought to test the hypothesis that these genes encode EcKs by misexpression, driving UAS-ORF constructs (and an Epgy2 insertion upstream of the native *Wall* locus that contains a UAS and promoter region—UAS-*Wall^EY^*) with the strong ubiquitous driver *tub-GAL4* at 25 °C to see if ectopic expression arrested developmental progression. As a positive control for ecdysteroid inactivation, we used a UAS-*Cyp18a1* responder (Guittard et al. 2011) to misexpress the ecdysteroid 26-hydroxylase/carboxylase Cyp18a1, which phenocopies ecdysteroid biosynthesis mutants (Guittard et al. 2011; Rewitz et al. 2010).

Ubiquitous misexpression of *Cyp18a1* with *tub-GAL4* completely arrested development, as expected, as did ubiquitous misexpression of *Wall* with either the UAS-*Wall^EY^* line or the UAS-*Wall^pU^* line. Examination of offspring vials under a fluorescent microscope (to detect the presence of the TM3 *actGFP Ser^1^* balancer chromosome) suggested that these misexpression genotypes failed to complete embryogenesis and hatch as larvae (Figure 3A). Misexpression of either *pkm* ORF transgene failed to significantly alter adult genotypic ratios from Mendelian expectations, suggesting ubiquitous expression of either *pkm* ORF does not affect developmental progression (Figure 3A).

**Figure 3.**
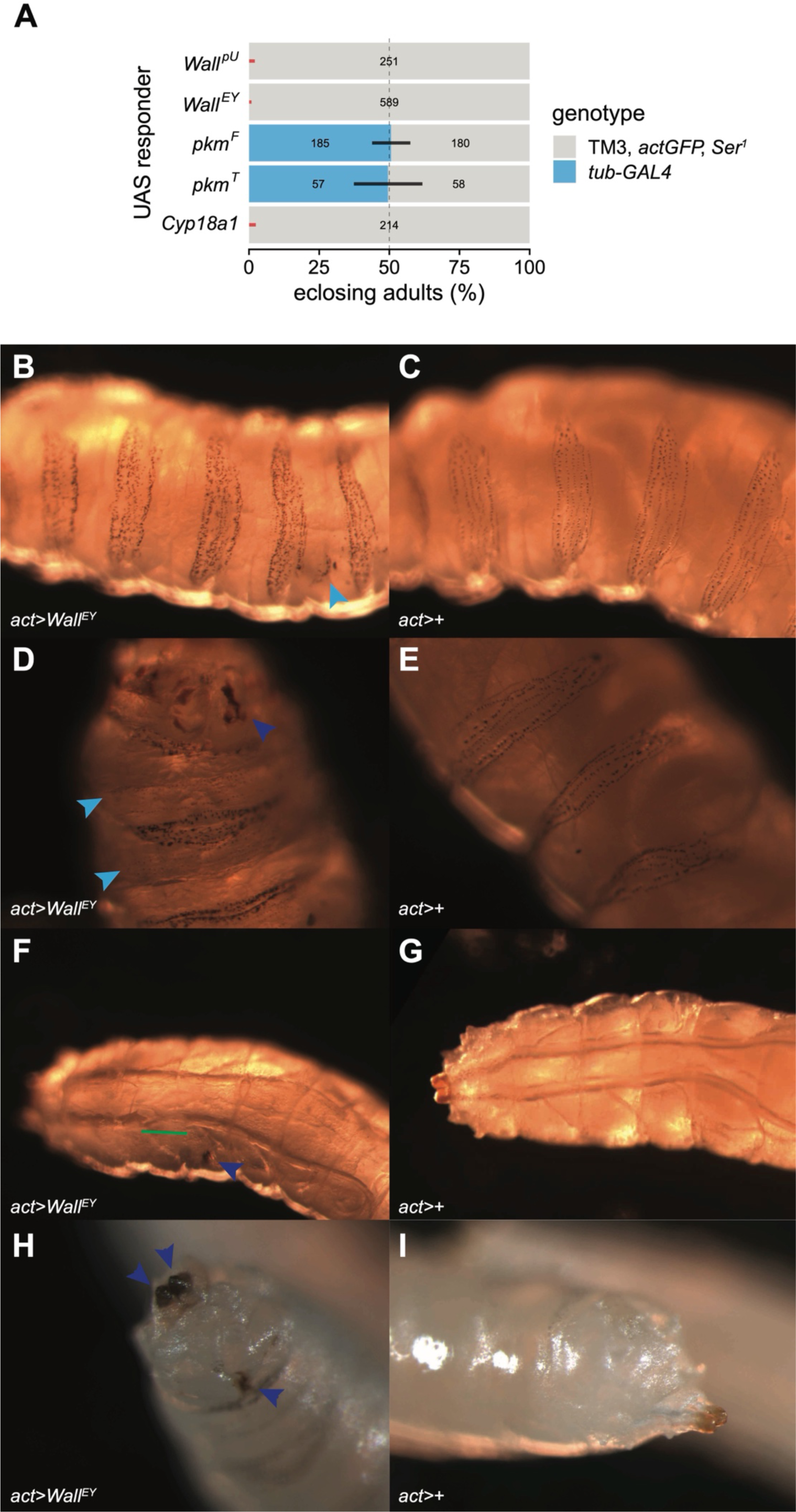
Ubiquitous misexpression of *Wall*, *pkm* and *Cyp18a1*. (A) Egg-to-adult viability of strong, ubiquitous *Wall*, *pkm* and *Cyp18a1* misexpression, estimated from the adult genotypic ratios of offspring from crosses between *tub-GAL4*/TM3 *actGFP Ser^1^* females (genotypes in bold) and UAS-responder males at 25 °C. UAS-*pkm^F^* expresses the full 635 aa *pkm* ORF, while UAS-*pkm^T^* expresses a truncated 430 aa ORF that does not include the pkm N-terminal disordered region. The dashed line indicates the expected 1:1 genotypic ratio if both genotypes per cross are equally developmentally viable; error bars are 99% confidence intervals (95% CI adjusted for five tests) for the proportion of *GAL4*-containing heterozygotes; black and red bars indicate non-significant or significant deviations, respectively, from expected genotypic ratios after correction for multiple tests. Numbers on the bars are the number of adults of each genotype (for numbers greater than 0). (B–I) Phenotypes in larvae misexpressing UAS-*Wall^EY^* with the *act-GAL4* driver (*act>Wall^EY^*; left) compared with wild-type, non-misexpressing larvae (*act>*+; right), at 25 °C. (B–E) Denticle belt disorganisation, including ectopic denticles in between belts (light blue arrowheads) and rectal pad melanisation (dark blue arrowheads). (F–G) Fluid-filled section of a tracheal dorsal trunk (green line segment), along with a melanised section of trachea (dark blue arrowhead). (H–I) Melanised posterior spiracles and rectal pad (dark blue arrowheads).

Misexpression of *Wall* using UAS-*Wall^EY^* and the strong, ubiquitous *act-GAL4* driver at 25 °C also caused complete developmental arrest (n = 158 adults, p < 2.2×10^-16^, exact binomial test), although not all animals died as embryos, with some larvae hatching but dying before pupariation, with various morphological phenotypes, such as spiracle and rectal pad pigmentation, and tracheal and denticle belt defects (Figure 3B–I).

### Ubiquitous misexpression of *Wall* causes developmental arrest until the middle of metamorphosis

While ubiquitous misexpression of *Wall* is embryonic lethal and occasionally larval lethal, it was unclear if misexpression affected later developmental stages. To investigate this, a delayed-onset misexpression experiment was conducted using the temperature-sensitive *tub-GAL80^ts^* construct, which represses GAL4 activity at 18 °C and permits GAL4 activity at 29 °C, and the *act-GAL4* driver, which strongly expresses GAL4 in a ubiquitous pattern (*tub-GAL4* was not used because pilot experiments suggested *tub-GAL80^ts^*was unable to fully repress its activity). Misexpression of *Wall* caused detectable drops in egg-to-adult viability when delayed 0–14 days into development, but did not affect viability from 15–18 days (at 18° C adult emerge at ∼18 days), suggesting animals become insensitive to *Wall* misexpression around the middle of metamorphosis (Figure 4).

**Figure 4.**
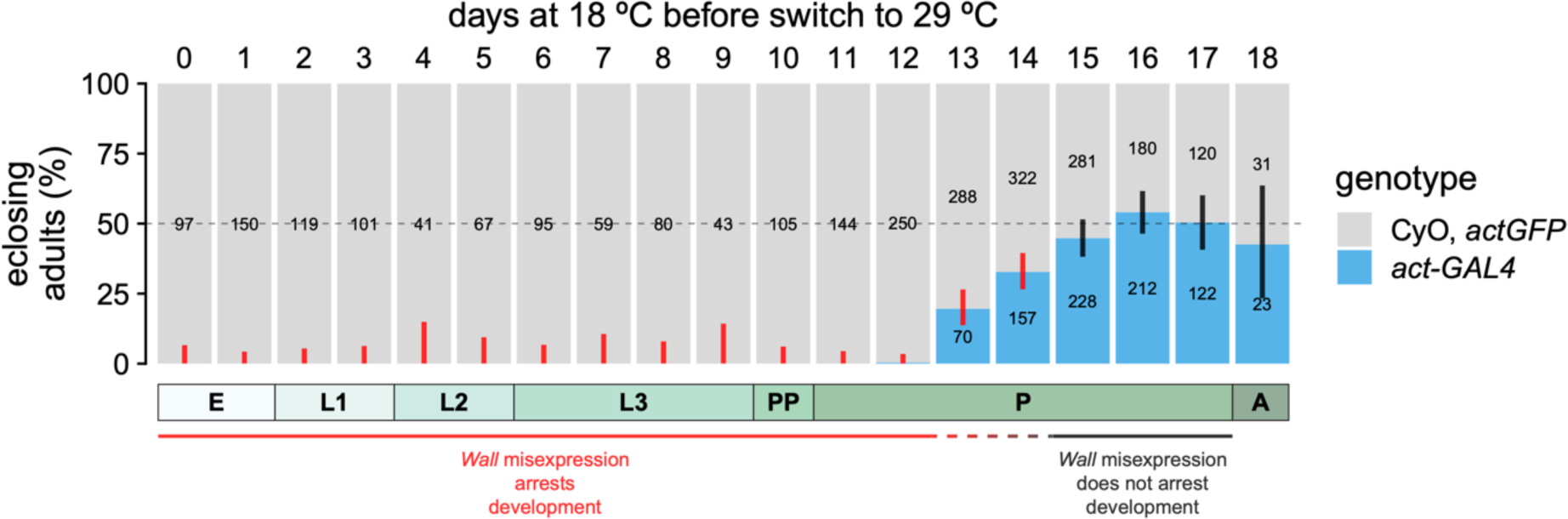
Delayed onset of ubiquitous misexpression of *Wall* arrests development up until mid-metamorphosis. Top: Egg-to-adult viability of *Wall* misexpression with the ubiquitous driver *act-GAL4*, in concert with the UAS-*Wall^EY^* element and the temperature-sensitive *tub-GAL80^ts^* construct, which inhibits GAL4 at 18 °C but not at 29 °C; animals were moved from 18 °C to 29 °C at one of 19 timepoints (top), which began *Wall* misexpression. Viability is estimated from the adult genotypic ratios of offspring from crosses between *act-GAL4*/CyO *actGFP* females and *tub-GAL80^ts^*; UAS-*Wall^EY^* males. The dashed line indicates the expected 1:1 genotypic ratio if both genotypes per cross are equally developmentally viable; error bars are 99.7% confidence intervals (95% CI adjusted for 19 tests) for the proportion of *act-GAL4*-containing heterozygotes; black and red bars indicate non-significant or significant deviations, respectively, from expected genotypic ratios after correction for multiple tests. Numbers on the bars are the number of adults of each genotype (for numbers greater than 10). Bottom: Approximate developmental life stages of *D. melanogaster* at 18 °C, showing *Wall*-sensitive (red) and *Wall*-insensitive (black) developmental periods; dashed line indicates region of uncertainty. E, embryo; L1, 1^st^-instar larva; L3, 2^nd^-instar larva; L3, 3^rd^-instar larva; PP, prepupa; P, pupa; A, adult.

### Tissue-specific misexpression of *Wall* causes developmental arrest distinct from *Cyp18a1*

The developmental arrest phenotype observed upon ubiquitous misexpression of *Wall* is consistent with an EcK function of its encoded protein. To further test this hypothesis, we conducted tissue-specific misexpression crosses to see if the patterns of developmental arrest produced by *Cyp18a1* misexpression matched that of *Wall* misexpression—a close match would be good evidence that Wall and Cyp18a1 act on the same or similar substrates.

Misexpression of *Wall* was conducted with two ubiquitous GAL4 drivers (*tub-GAL4*, used previously, and *da-GAL4*, a weaker driver) and 14 tissue-specific GAL4 drivers (Table S1). The positive control UAS-*Cyp18a1* construct produced developmental arrest when misexpressed with all 16 drivers (Figure 5; Table S4). The UAS-*Wall^EY^* construct produced developmental arrest when misexpressed with 10 drivers, while the UAS-*Wall^pU^*construct produced very similar results, but with no phenotype with the *ppl-GAL4* driver and generally caused developmental arrest at later life stages; *btl-GAL4* and *cg-GAL4* misexpression also resulted in a small number of adult escapers (Figure 5, Table S4). These differences are likely due to differences in expression strength: the *EPgy2* element contains 14 copies of UAS in its artificial enhancer region (Bellen et al. 2004), while pUASTattB contains only five (Bischof et al. 2007). However, the broad consistency of the results from both an independent transgene construct and a native locus misexpression construct means these phenotypes can be confidently linked to the ectopic expression of the *Wall* ORF.

**Figure 5.**
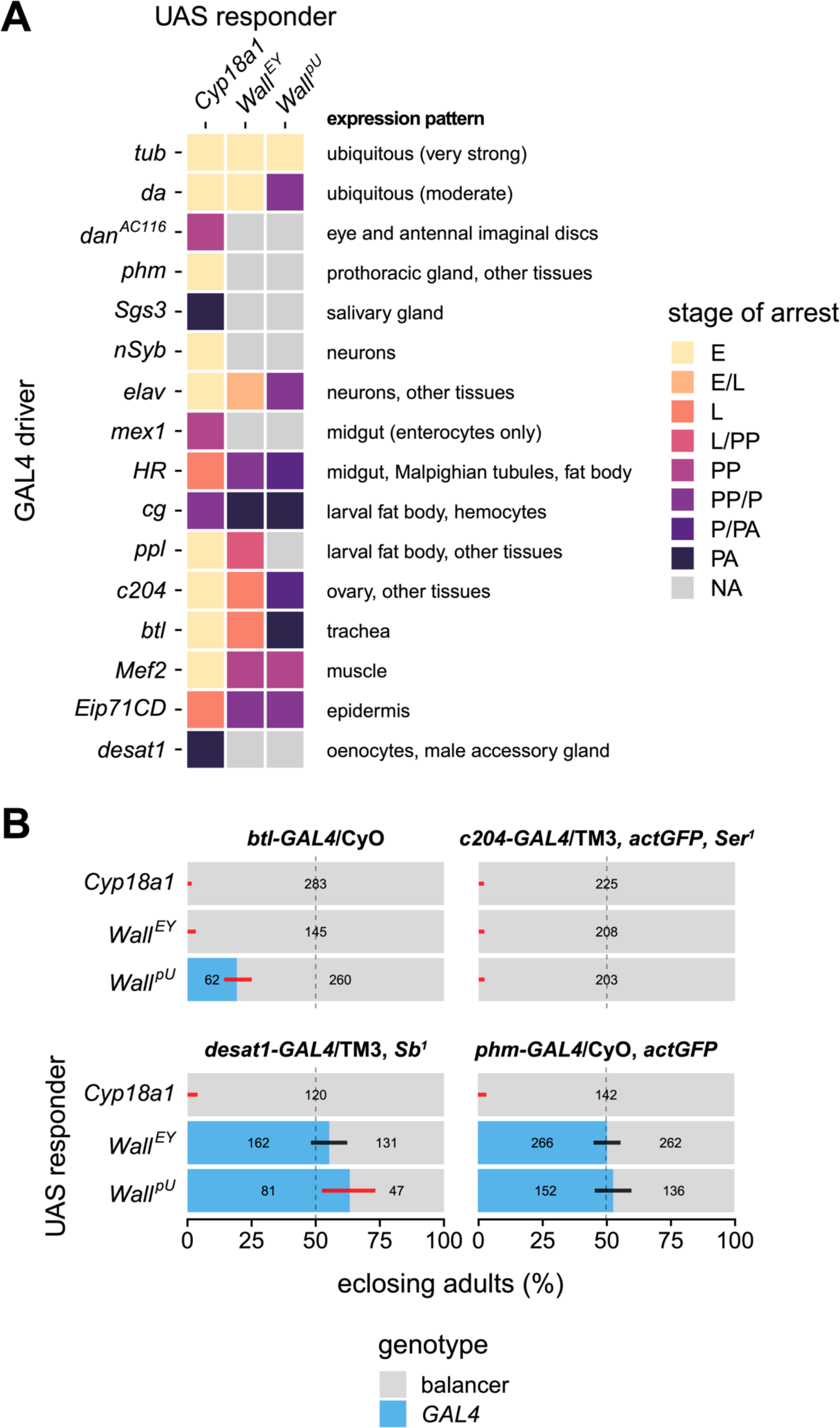
Tissue-specific misexpression of *Wall* and *Cyp18a1*, using the UAS-*Wall^EY^*, UAS-*Wall^pU^* and UAS-*Cyp18a1* constructs, at 25 °C. (A) Heat map of qualitative developmental arrest phenotypes for *Wall* and *Cyp18a1* misexpression using two ubiquitous GAL4 drivers and 14 tissue-specific GAL4 drivers. E, embryo; L, larva; PP, prepupa; P, pupa; PA, pharate adult; NA, no arrest (successful eclosion). When arrest occurred at more than one stage, two stages are indicated and separated with a slash. For phenotypic descriptions, see Table S4. (B) Egg-to-adult viability of *Wall* and *Cyp18a1* misexpression, estimated from the adult genotypic ratios of offspring from crosses between *GAL4*/balancer females (genotypes in bold) and UAS-responder males. The dashed line indicates the expected 1:1 genotypic ratio if both genotypes per cross are equally developmentally viable; error bars are 98.3% confidence intervals (95% CI adjusted for three tests) for the proportion of *GAL4*-containing heterozygotes; black and red bars indicate non-significant or significant deviations, respectively, from expected genotypic ratios after correction for multiple tests. Numbers on the bars are the number of adults of each genotype (for numbers greater than 0).

Misexpressing UAS-*Wall^EY^* with the *HR-GAL4* driver resulted in a dramatic phenotype: an extended wandering period during the 3^rd^ larval instar, then defects during pupariation and pupation wherein the puparium was thin and elongated and anterior adult tissue differentiation appeared to occur without head eversion, resulting in dark thoracic (likely from wings) and red eye pigmentation developing internally (Figure S6A–C). This phenotype suggests *Wall* misexpression with this driver can block epidermal and cuticular development during metamorphosis, but curiously allows for imaginal discs eversion to take place. The *HR-GAL4* driver expresses GAL4 along the whole length of the midgut, as well as in the Malpighian tubules and the fat body (Chung et al. 2007); however, the lack of phenotypes with the enterocyte-specific *mex1-GAL4* driver suggests that *Wall* misexpression in these midgut cells does not cause developmental defects (Phillips & Thomas 2006). Additionally, given that the fat body drivers *cg-GAL4* and *ppl-GAL4* failed to phenocopy *HR-GAL4*, the crucial tissues might be non-enterocyte midgut cells and/or the Malpighian tubules. Alternatively, *HR-GAL4* may drive at a low (yet sufficient) level in another tissue to produce this phenotype, such as the muscles.

Conspicuously, while the *phm-GAL4* driver, which expresses GAL4 in the prothoracic cells of the ring gland (PG)—the ecdysteroidogenic tissue in larvae (Redfern 1983)—as well as some other tissues such as the wing disc (Casas-Tintó et al. 2017), caused embryonic lethality when misexpressing *Cyp18a1* as expected, yet it produced no developmental arrest when misexpressing either *Wall* construct (Figure 5B). This result appears strongly inconsistent with *Wall* encoding a kinase that can act on ecdysteroids present in the PG.

Misexpression of *Wall* with the *c204-GAL4* driver caused larval arrest with the UAS-*Wall^EY^* construct and arrest during metamorphosis with the UAS-*Wall^pU^* construct (Figure 5). The expression pattern of this driver is not well characterised—while it is known to be expressed in the adult ovary (Manseau et al. 1997), it may be expressed in other tissues and at other life stages, given it causes embryonic arrest with UAS-*Cyp18a1* and the aforementioned arrest with UAS-*Wall* constructs. Curiously, *elav-GAL4* misexpression of *Wall* caused developmental arrest, while *nSyb-GAL4* misexpression did not, suggesting misexpression in neurons is not the cause of *elav-GAL4* arrest; the *elav-GAL4* driver, while often considered specific to neurons, expresses in other tissues, such as glia, trachea and the imaginal discs (Berger et al. 2007; Casas-Tintó et al. 2017), one or more of which is likely the cause of the developmental arrest observed here.

Overall, these data demonstrate that *Wall* misexpression in the trachea, muscle, epidermis or fat body is sufficient to cause developmental arrest in *D. melanogaster*, while misexpression in the PG, imaginal discs, salivary gland, neurons and oenocytes does not cause any obvious phenotype. In addition, combined with the results of the ubiquitous delayed-onset misexpression experiment (Figure 4), these data suggest that the pharate adult lethality observed through misexpression with some drivers (*cg-GAL4* and *btl-GAL4*) is likely due to misexpression of *Wall* during early metamorphosis or earlier, not during the pharate adult stage itself.

### Larval tracheal misexpression of *Wall* causes tracheal defects and food aversion

Misexpression of *Wall* with the trachea-specific *btl-GAL4* driver caused developmental arrest at the larval or pharate adult stages, depending on the UAS-responder construct and the temperature (Figure 5, Table S4). We noticed that *btl>Wall^EY^* larvae raised at 25 °C abandoned the food media to crawl up the sides of the vial, a phenotype that was shared with ubiquitously misexpressing *da>Wall^EY^* larvae that were escaped from embryonic lethality using the *tub-GAL80^ts^* construct and then moved to 29 °C, as well as the *act>Wall^EY^* larvae described previously.

Quantification of this food-aversion phenotype showed *da>Wall^EY^*larvae were positioned differently within vials compared to non-misexpressing *da>w^1118^* controls, at both two days and three days after hatching (both p < 2.2×10^-16^, Fisher’s exact test; Figure 6A). There was also a significant difference in positions between the two timepoints for misexpressing larvae (p = 9.7×10^-14^, Fisher’s exact test), while there was no significant difference between non-misexpressing larvae (p = 0.73, Fisher’s exact test), consistent with progression of the misexpression phenotype over time (Figure 6A). Food aversion is a well-established consequence of hypoxia in both larvae and adult *D. melanogaster* (Wang et al. 2015; Vigne & Frelin 2010; Wingen et al. 2017)—consistent with this, close examination of *btl>Wall^EY^* and *da>Wall^EY^* larvae revealed defects in tracheal filling (similar to those seen in *act>Wall^EY^*and *c204>Wall^EY^* larvae; Figure 3F, Table S4), with parts of the dorsal trunks often filled with liquid (Figure 6B), which can cause severe reductions in gas exchange (Parvy et al. 2012). Taken together, these data suggest that *Wall* misexpression in trachea causes hypoxia through defects in either tracheal development, maintenance or integrity. The food-aversion phenotype is also the basis for the gene name *Wallflower* (*Wall*), due to the tendency of the larvae to position themselves on the walls of the vial, rather than in the food with other larvae, an allusion to the ‘wallflower’ metaphor for social behaviour.

**Figure 6.**
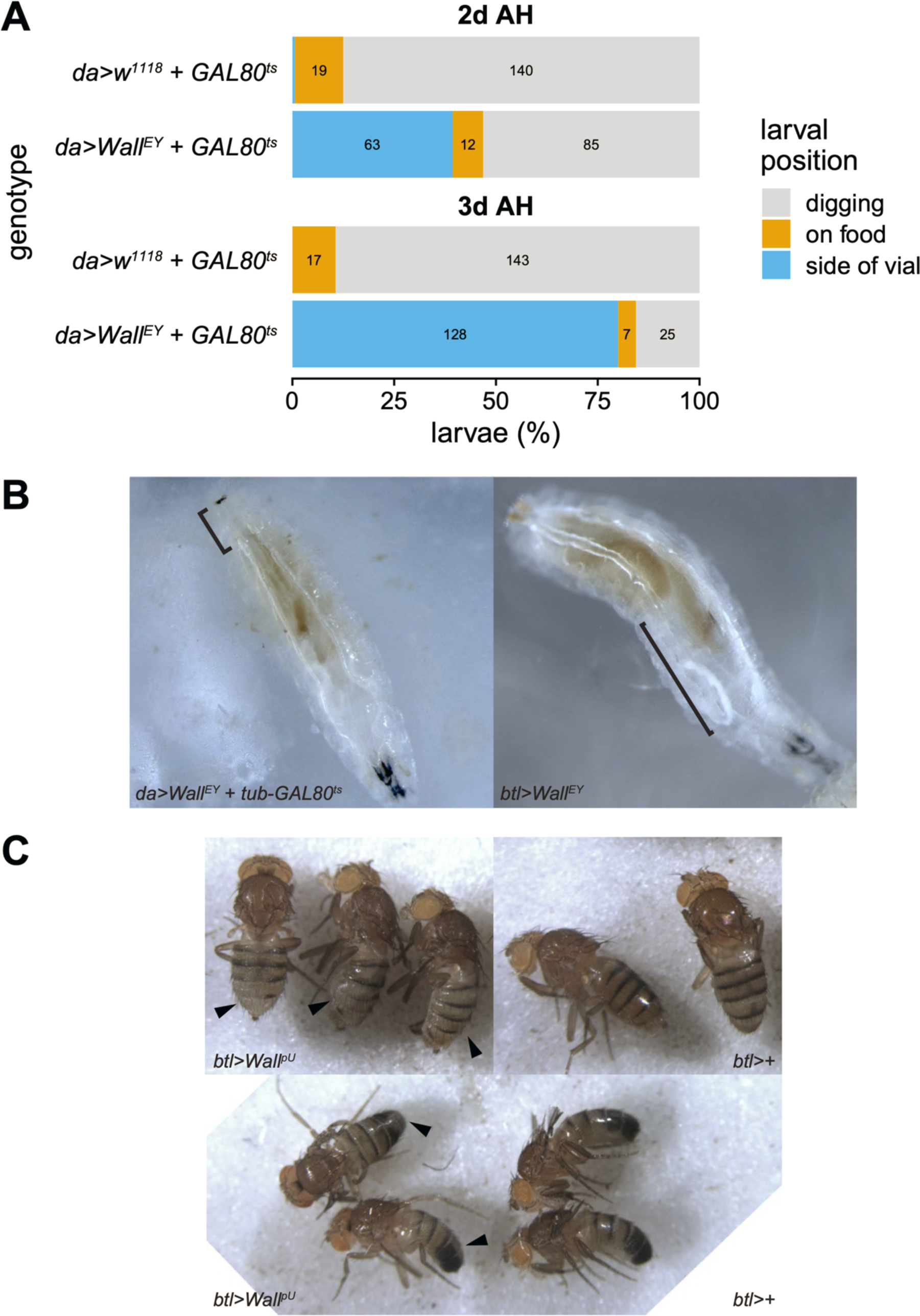
Misexpression of *Wall* affects tracheal integrity and adult pigmentation. (A) Food avoidance assay comparing *da>w^1118^ + GAL80^ts^* and *da>Wall^EY^* + *GAL80^ts^* larvae 2 days (top) and 3 days (bottom) after hatching (AH). Numbers on the bars are the number of larvae in each position (for numbers greater than five). (B) Photos of larvae misexpressing *Wall* ubiquitously (*da>Wall^EY^* + *tub-GAL80^ts^*, left) or specially in the trachea (*btl>Wall^EY^*, right) retrieved from food media at 25 °C, before food aversion occurred. Black segments indicate sections of tracheal dorsal trunks that are filled with liquid. (C) Adult flies (females, top; males, bottom) misexpressing *Wall* in the trachea (*btl>Wall^pU^*, left) or with the *btl-GAL4* driver alone (*btl>+*, right). Arrowheads indicate dorsal abdomen pigmentation defects (areas of lightened pigment compared to the rest of the abdomen). Wings have been removed to expose the dorsal abdomen.

Misexpression of *Wall* with the *btl* driver also appears to affect tissues other than the trachea, as we noticed a pigmentation defect on the dorsal abdomen of both male and female *btl>Wall^pU^* adult escapers (Figure 6C).

### *Wall* misexpression phenotypes are not hypostatic to *Cyp18a1*

We hypothesised that *Wall* encodes an ecdysteroid 26-kinase, which must act on the products of the ecdysteroid 26-hydroxylase Cyp18a1. If this hypothesis is true, *Wall* misexpression should be hypostatic to the wild-type function of *Cyp18a1*—ie. misexpression of an ecdysteroid 26-kinase (Wall) should not cause developmental defects when the source of 26-hydroxyecdysteroids (Cyp18a1) is not functional. Evidence of hypostasis would be strong genetic evidence of a biochemical interaction between the two genes and would support the hypothesis that *Wall* encodes an ecdysteroid 26-kinase. In contrast to *Cyp18a1^1^* animals, which largely reach the pharate adult stage but fail to eclose (Rewitz et al. 2010), most *da>Wall^pU^* animals at 25 °C arrest development before the differentiation of adult structures (prepupal/pupal arrest), while most *Mef2>Wall^pU^* animals arrest development before pupation (prepupal arrest). We reasoned that if *Wall* misexpression phenotypes were hypostatic to the function of *Cyp18a1*, *Cyp18a1^1^* hemizygotes (25% of the offspring of crosses between *Cyp18a1^1^*/FM7i, *actGFP*; UAS-*Wall^pU^* females and GAL4 males) should reach the pharate adult stage instead of earlier developmental arrest due to *Wall* misexpression, increasing the proportion of pharate adult-lethal animals.

There were no significant differences in developmental outcomes between *Cyp18a1^1^*, *Mef2>Wall^pU^* and *Mef2>Wall^pU^*offspring (Figure S7A; p = 0.336, Fisher’s exact test), nor between *Cyp18a1^1^*, *da>Wall^pU^* and *da>Wall^pU^* offspring (Figure S7B; p = 0.841, Fisher’s exact test). These data indicate that misexpression of *Wall* in *Cyp18a1^1^* hemizygotes produces the same developmental arrest phenotypes as misexpression of *Wall* in a wild-type background, strongly suggesting that *Wall* is not hypostatic to *Cyp18a1*, and that *Wall* is unlikely to encode an ecdysteroid 26-kinase.

### *pkm* mutants are not sensitive to starvation

*pkm* was found by Chatterjee et al. (2014) to be up-regulated in adult female oenocytes upon starvation, which led to the hypothesis that *pkm* plays a role in the metabolic response to starvation, and that *pkm* loss-of-function mutants are more susceptible to starvation. Wet-starvation experiments (Storelli et al. 2019) were conducted with virgin male and virgin female flies, the offspring from three different crosses: *pkm^4^* females and *w^1118^* males (yielding *pkm^4^*/*pkm^+^* females and *pkm^4^*/Y males); *pkm^4^* females and *pkm^2^* males (yielding *pkm^4^*/*pkm^2^* females and *pkm^4^*/Y males); and *w^1118^* females and *pkm^2^* males (yielding *pkm^2^*/*pkm^+^* females and *pkm^+^*/Y males). For heterozygous female genotypes (Figure S8A), median survival in days were 9 (95% CI: 8, 9), 8 (95% CI: 8, 8) and 9 (95% CI: 8, 9) for *pkm^4^*/*pkm^2^*, *pkm^4^*/*pkm^+^* and *pkm^2^*/*pkm^+^*, respectively, with no significant differences between any pairs of genotypes across entire survival curves (all p > 0.6, log-rank test). For hemizygous male genotypes (Figure S8B), median survival in days were 7 (95% CI: 6, 7), 8 (95% CI: 7, 9) and 6 (95% CI: 6, 7) for *pkm^4^*/Y (*pkm^2^*father), *pkm^4^*/Y (*w^1118^* father) and *pkm^+^*/Y (*pkm^2^* father), respectively, with the only significant difference between the *pkm^4^*/Y (*pkm^2^* father) and *pkm^4^*/Y (*w^1118^* father) genotypes (p = 0.0037, log-rank test; all other p > 0.067) across entire survival curves. These data do not support the hypothesis that *pkm* is involved in starvation resistance.

### Misexpression of *CG5644*, another strong EcK candidate gene, also causes developmental arrest

As other EcKLs in the *D. melanogaster* genome may also encode EcKs, we screened an additional 13 EcKLs for misexpression phenotypes using pre-existing UAS-ORF fly lines and the *tub-GAL4* driver. Six of these genes—*CG31300*, *CG31288*, *CG10560*, *CG10562*, *CG6830* and *JhI-26*—were previously considered candidate detoxification genes, while the seven others— *CG10514*, *CG31102*, *CG31087*, *CHKov2*, *CG9259*, *CG14314* and *CG5644*—were not (Scanlan et al. 2020).

Only misexpression of *CG6830* and *CG5644* caused significant developmental arrest before the adult stage (Figure S10A). The UAS-*CG5644* line was further crossed to the *phm-GAL4* driver at 25 °C, resulting in 66 misexpression adults surviving compared to 115 balancer adults (proportion = 0.365, 95% CI: 0.295– 0.439, p = 0.0003, binomial test), suggesting PG misexpression of *CG5644* can also disrupt development. These data are consistent with the hypothesis that *CG5644* encodes an ecdysteroid 22-kinase, previously proposed based on 1:1 orthology to *AgEcK2* in *Anopheles gambiae* (Peng et al. 2022; Scanlan & Robin 2023). The lack of developmental arrest seen in *tub>CG31300* and *tub>JhI-26* individuals is consistent with previously published experimental data (Liu et al. 2014; Scanlan et al. 2022).

While misexpression of *CG31288* did not appear to affect developmental progression, we noticed a visible phenotype for *tub>CG31288* adult flies: their meconium was a dark grey, almost black colour, compared to the yellow-green of non-misexpressing flies (Figure S10B–D). This colour change was apparent both within the posterior abdomen of recently eclosed flies (Figure S10B) and when the meconium had been excreted and left on empty puparia (Figure S10C–D).

## Discussion

Here we have performed the first detailed genetic characterisation of candidate ecdysteroid kinase (EcK) genes in insects, targeting *Wall* and *pkm*, orthologs of the known ecdysteroid 22-kinase gene *BmEc22K* in the silkworm *B. mori*. Despite phenotypic discrepancies between gene disruption methods, we ultimately found no clear evidence that these genes encode EcKs that have essential functions in development or reproduction. Tissue-specific misexpression was also used to detect possible EcK activity, which produced striking developmental disruption phenotypes for *Wall* but not for *pkm*.

The phenotypic differences we observed between our null allele, RNAi knockdown and somatic CRISPR knockout approaches were unexpected. Broadly, strong RNAi knockdown conditions resulted in developmental arrest for both genes, across independent dsRNA constructs, but homozygous mutant animals (including double mutants) showed no obvious developmental phenotypes and had similar viability to controls (see Table S3 for a summary). One explanation is genetic compensation in null mutants that isn’t trigged during RNAi knockdown (El-Brolosy & Stainier 2017), but knocking down *pkm* in a *pkm* mutant background did not rescue developmental arrest, inconsistent with this hypothesis. The more likely explanation then is that the RNAi knockdown phenotypes were due to off-target effects, which have been well documented in *Drosophila* (Seinen et al. 2011). For *Wall*, no phenotypes were detected for both null alleles and somatic CRISPR knockout, however for *pkm*, somatic CRISPR knockout produced a highly penetrant motor coordination defect not seen in null mutant animals. Explanations for this are unclear, but could be linked to CRISPR-Cas9 off-target cleavage or unintended effects on the splicing or transcriptional regulation of *nod*, the fourth intron of which *pkm* sits inside.

Null alleles are generally considered the gold standard for characterisation of gene function. Animals homozygous or hemizygous for null alleles of *Wall*, *pkm* or both genes appeared indistinguishable from wild-type animals respect to both developmental progression and reproduction, strongly suggesting these two closely related EcKLs do not play important roles in these processes, even redundantly with each other. This is in contrast with *BmEc22K*, another subfamily A EcKL that encodes an ecdysteroid 22-kinase involved in ovary–embryo ecdysteroid recycling (Sonobe et al. 2006), and indicates that this function is not conserved in dipteran subfamily A genes. It is possible *Wall* and/or *pkm* null mutants have subtle or environmentally specific phenotypes we were unable to detect in our experiments, although the lack of expression of either gene in reproductive tissues (Leader et al. 2018) makes this unlikely for reproductive functions. The lack of strict conservation of the EcKL clades containing *pkm* and *Wall* (referred to as Dip9 and Dip10 in Scanlan & Robin 2023) is consistent with non-essential functions of these genes in *Drosophila* and other dipteran taxa.

Neither null mutants nor misexpression animals for *pkm* displayed any obvious phenotypes, offering no clue as to the gene’s native function or enzymatic activity. We hypothesised that *pkm* is involved in the starvation response in adult flies, based on transcriptomic data from Chatterjee *et al*. (2014), but experiments with null mutants did not show a difference in wet starvation survival compared to control genotypes, for either sex, offering no support for the hypothesis. Using RNAi knockdown with the same dsRNA lines used here, Santana *et al*. (2020) suggested the Pkm protein controls mRNA and protein abundances of Hsp23 and Hsp26, leading to modulation of synaptogenesis and neural response to stress. However, our results cast doubt on the interpretability of RNAi knockdown of *pkm* and suggest off-target effects could be responsible for these phenotypes. Confirmation of neural functions of *pkm* could be done with null alleles, which we show here to be viable and therefore could allow the study of subtle neurological phenotypes at any life stage.

Like *pkm*, *Wall* null mutants had no obvious phenotypes. However, misexpression of *Wall* disrupted developmental progression with both ubiquitous and tissue-specific drivers, suggesting the Wall protein can act on one or more developmentally important small molecules. Tissues sensitive to *Wall* misexpression are the trachea, fat body, epidermis and muscle, while insensitive tissues are the PG, neurons, oenocytes, imaginal discs and salivary glands. Ubiquitous misexpression of *Wall* caused developmental arrest at all life stages, except past the midpoint of metamorphosis, after pupal–adult apolysis has occurred (Bainbridge & Bownes 1981). These data suggest the substrate(s) of Wall could have essential signalling functions in trachea, fat body, epidermis and muscle across *D. melanogaster* development. However, it is unlikely these substrates are E and/or 20E, as: 1) embryonic *Wall* misexpression did not phenocopy the “Halloween” phenotype like *Cyp18a1* (Rewitz et al. 2010); 2) *Wall* phenotypes were specific to certain tissues, while *Cyp18a1* misexpression reliably caused developmental arrest in tissues known to rely on 20E signalling; and 3) *Wall* misexpression in the PG, the larval source of E, did not produce developmental arrest, unlike *Cyp18a1*. Curiously, however, a recent paper reports that the dorsal internal oblique muscles (DIOMs) are the source of the mid-pupal pulse of 20E (Zhang et al. 2023); metamorphic defects seen upon pan-muscular *Wall* misexpression could be consistent with interfering with ecdysteroid signalling in DIOMs, although these generally occurred before the middle of metamorphosis (Figure 5A).

An ecdysteroid 26-kinase is known to be active in the S2 cell line (Guittard et al. 2011), which we hypothesised could be encoded by *Wall* due to transcriptional similarities to the ecdysteroid 26-hydroxylase/carboxylase gene *Cyp18a1*. However, this hypothesis was not supported by epistasis experiments, which found that the severity of *Wall* misexpression phenotypes was not reduced in a *Cyp18a1* mutant background, suggesting *Wall*-induced developmental defects are to not due to phosphorylation of 26-hydroxylated ecdysteroids produced by Cyp18a1. Alternative candidates for encoding this ecdysteroid 26-kinase are *CG2004*, *CG31975* and *JhI-26*, which are all basally expressed in S2 cells to appreciable degrees (Figure S10). Plausible, alternative ecdysteroid substrates for Wall are 3-dehydroecdysteroids and 3-epiecdysteroids, which are produced in the fat body and epidermis of larvae (Sommé-Martin et al. 1988a, 1988b) and may have signalling functions in *D. melanogaster* (Sommé-Martin et al. 1990; Baker et al. 2003). The physiological functions of these modified ecdysteroids are still unclear in *D. melanogaster*, but intriguingly, 3-dehydro-20E was recently found to modulate female reproductive physiology in the mosquito *Anopheles gambiae* (Peng et al. 2022). Additionally, Cavigliasso *et al*. (2023) recently suggested *fezzik* (*fiz*) encodes an ecdysteroid oxidase that adaptively modulates larval growth rate under malnutrition in *D. melanogaster* (Grenier et al. 2023; Glaser-Schmitt & Parsch 2018). If oxidised ecdysteroids control larval development, they would be good candidate substrates for Wall. Other possible substrates could be produced by Cyp301a1, Cyp303a1 and Spidey, developmentally important yet biochemically uncharacterised enzymes with various links to ecdysteroid metabolism (Sztal et al. 2012; Wu et al. 2019; Chiang et al. 2016).

The phenotypes associated with *Wall* misexpression might not necessarily reflect the gene’s wild-type function, as the enzyme may be acting promiscuously on substrates that it rarely or never encounters in non-transgenic animals. However, if they are attributable to native substrates, these misexpression phenotypes might still be exposing novel hormonal pathways that underpin important developmental processes. Such native substrates would clearly be involved in tracheal integrity, cuticle and epidermal development, and functions of the fat body and muscles during metamorphosis. These are all known to be regulated by ecdysteroid signalling (Buhler et al. 2018; Bond et al. 2010; Zachary & Hoffmann 1980; Chanut-Delalande et al. 2014), but links to signalling genes and proteins that have been associated with more than one of the developmental defects seen in *Wall*-misexpressing animals could offer mechanistic clues. These include: the scavenger receptor Dsp, which is involved in both tracheal integrity and adult pigmentation (Dembeck et al. 2015; Wingen et al. 2017); *Megf8*, which is involved in tracheal integrity and denticle belt development (Lloyd et al. 2018); and Svb, which has roles in tracheal and denticle belt development (Chanut-Delalande et al. 2006, 2014). Intriguingly, Svb also acts in the adult midgut to coordinate stem cell behaviour under the control of EcR (Al Hayek et al. 2019), which is a major site of native *Wall* expression (Leader et al. 2018). Ultimately, biochemical characterisation of the purified Wall protein is needed to fully understand its signalling interactions upon misexpression.

The results of this study leave as open questions the identities of the ecdysteroid 2-, 3-, 22- and 26-kinases in *D. melanogaster*. However, data from our misexpression experiments is consistent with *CG5644* encoding an EcK, especially the developmental arrest seen upon PG-specific misexpression of this gene. Our previous phylogenetic analyses of the EcKL gene family determined that *CG5644* is a 1:1 ortholog of *AgEcK2*, which encodes an ecdysteroid 22-kinase in *A. gambiae* (Peng et al. 2022; Scanlan & Robin 2023). While *CG5644* is predominantly expressed in the adult nervous system (Leader et al. 2018), it also appears expressed in the migratory border cells in the ovary (Borghese et al. 2006), which might make it responsible for the presence of E 22-phosphate in this tissue (Pis et al. 1995; Grau et al. 1995). Reverse genetics experiments like those conducted here for *Wall* and *pkm* and biochemical characterisation might uncover the functions of both *CG5644* and ecdysteroid 22-phosphates in *D. melanogaster* development, reproduction and physiology.

Overall, we here provide the first in-depth functional genetic characterisation of candidate EcK genes in insects, focusing on the *BmEc22K* orthologs *Wall* and *pkm* in *D. melanogaster*. While neither gene is essential for reproduction and development, multiple lines of evidence are consistent with Wall being able to inactivate ecdysteroid-like molecules. This work is suggestive of unknown complexity in tissue-specific hormonal signalling in *D. melanogaster* and paves the way for further characterisation of the likely ecdysteroid 22-kinase gene *CG5644*. This work also serves to highlight the possible phenotypic inconsistencies that may arise between functional genetic methodologies and reinforces the importance of multiple, complementary gene disruption approaches when characterising genes of unknown function.

## Acknowledgments

The authors would like to thank Simon Bullock for the pCFD6 plasmid, the Drosophila Genomics Resource Center for the UAS-ORF plasmids, and Philip Batterham, Michael Murray and Michael O’Connor for Drosophila stocks.

